# MetaMap: An atlas of metatranscriptomic reads in human disease-related RNA-seq data

**DOI:** 10.1101/269092

**Authors:** LM Simon, S Karg, AJ Westermann, M Engel, AHA Elbehery, B Hense, M Heinig, L Deng, FJ Theis

**Affiliations:** Helmholtz Zentrum München, German Research Center for Environmental Health, Institute of Computational Biology, Neuherberg, Germany; Institute for Molecular Infection Biology, University Würzburg, Würzburg, Germany; Helmholtz Institute for RNA-Based Infection Research (HIRI), Würzburg, Germany; Helmholtz Zentrum München, German Research Center for Environmental Health, Scientific Computing Research Unit, Neuherberg, Germany; Helmholtz Zentrum München, German Research Center for Environmental Health, Institute of Virology, Neuherberg, Germany; Department of Mathematics, Technische Universität München, Munich, Germany

**Keywords:** High performance computing, big data, RNA-seq, sequence read archive, metatranscriptomics, microbiome, virome, human disease, infection

## Abstract

**Background:** With the advent of the age of big data in bioinformatics, large volumes of data and high performance computing power enable researchers to perform re-analyses of publicly available datasets at an unprecedented scale. Ever more studies imply the microbiome in both normal human physiology and a wide range of diseases. RNA sequencing technology (RNA-seq) is commonly used to infer global eukaryotic gene expression patterns under defined conditions, including human disease-related contexts, but its generic nature also enables the detection of microbial and viral transcripts.

**Findings:** We developed a bioinformatic pipeline to screen existing human RNA-seq datasets for the presence of microbial and viral reads by re-inspecting the non-human-mapping read fraction. We validated this approach by recapitulating outcomes from 6 independent controlled infection experiments of cell line models and comparison with an alternative metatranscriptomic mapping strategy. We then applied the pipeline to close to 150 terabytes of publicly available raw RNA-seq data from >17,000 samples from >400 studies relevant to human disease using state-of-the-art high performance computing systems. The resulting data of this large-scale re-analysis are made available in the presented MetaMap resource.

**Conclusions:** Our results demonstrate that common human RNA-seq data, including those archived in public repositories, might contain valuable information to correlate microbial and viral detection patterns with diverse diseases. The presented MetaMap database thus provides a rich resource for hypothesis generation towards the role of the microbiome in human disease.

## Data Description

### Context

Recent studies have demonstrated the paramount importance of the microbiome for human health and disease [1]. For example, imbalance of the human gut microbiome was linked to non-communicable diseases such as obesity [2,3], diabetes [4], cardiovascular disease [5], chronic obstructive pulmonary disease [6], or colorectal carcinoma [7,8], to name just a few.

The advent of high-throughput sequencing technologies has revolutionized the life sciences. RNA-seq technology produces one of the most frequent next generation sequencing data types and has been applied to study a large number of biological samples relevant to human disease. The majority of the underlying raw data is freely accessible from data repositories such as the Gene Expression Omnibus (GEO) (>1,700 human RNA-seq data sets as of january 2018) or the Sequence Read Archive (SRA) [9].

However, these data are typically exclusively used for single species (i.e. human) transcriptomics such as differential gene expression or alternative splicing analysis [9,10]. Reads that do not map onto the human genome are considered noise or contamination and therefore generally ignored [11,12] (collectively about 9% of total reads, Fig. 1). Five years ago, it was postulated that interspecies interactions might be studied by simultaneous detection and quantification of RNA transcripts from a given host and a microbe via ‘dual’ RNA-seq [13]. Meanwhile this approach has been successfully applied to the interaction of mammalian cells with diverse bacterial [14] and viral pathogens [15–19].

**Figure 1.**
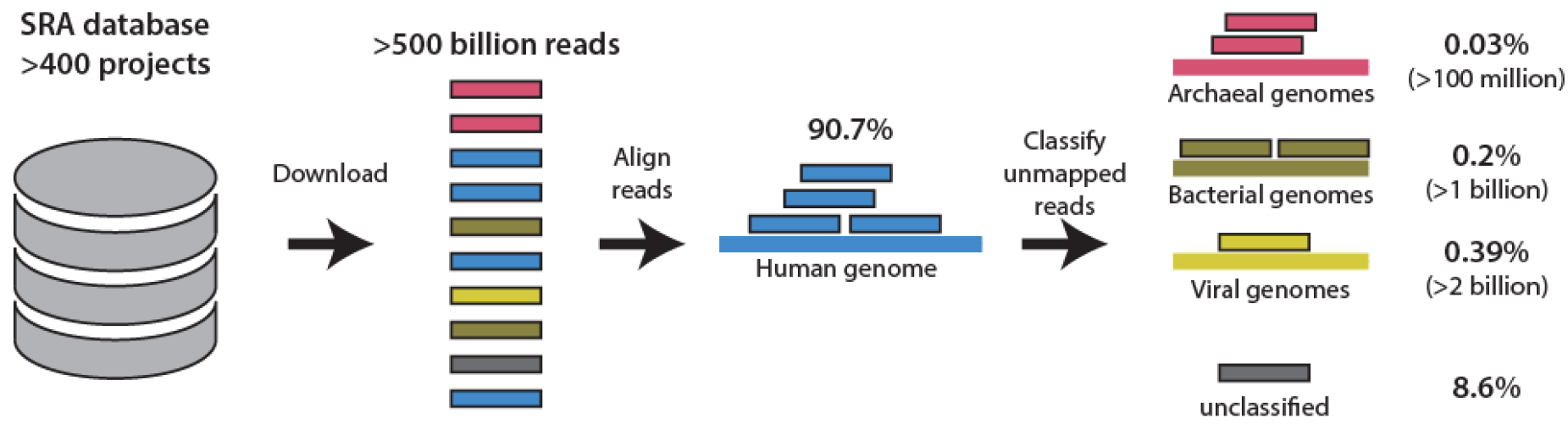
Schematic illustrates the MetaMap pipeline. Over 400 projects from studies relevant to human disease were identified in the SRA database. Over 500 billion RNA-seq reads were downloaded and first filtered by mapping them onto the human genome and subsequently the remaining reads underwent metafeature classification. 90.7% of all reads mapped to the human genome. 0.03%, 0.20% and 0.39% of all reads were assigned to archaeal, bacterial or viral metafeatures, respectively. 8.6% of all reads remain non-discriminative at the species level (‘unclassified’).

Inspired by dual RNA-seq, in this study we hypothesize that reads in archived RNA-seq datasets derived from human primary cells or tissue samples that fail to map against the human reference genome may contain valuable information about the presence of certain microbes in the respective body niches and/or under defined disease conditions. To enable metatranscriptomic study of these data, we combined existing read alignment and metagenomic classification software into a two-step ‘omni’ RNA-seq pipeline to comprehensively quantify archaeal, bacterial and viral reads in human RNA-seq data (Fig.1).

In the first step of this so called ‘Metamap’ pipeline, all reads are aligned against the human genome using the ultra-fast RNA-seq aligner STAR [20] and subsequently only the fraction of unmapped reads is subjected to metatranscriptomic classification using CLARK-S [21] (see Methods for details). The combination between scalability and accuracy was the main motivation behind choosing these two software packages over competing methods [22,23]. It is important to note that CLARK-S uses a set of uniquely discriminative short sequences at the species level to classify reads. Therefore, reads containing non-discriminative sequences that fail to be uniquely assigned to a single species, e.g. reads originating from the bacterial ribosomal 16S rRNA gene, will be considered ‘unclassified’ (altogether 8.6% in Fig. 1).

The output of CLARK-S is an operational taxonomic units (OTU) count matrix, where rows correspond to viral, bacterial and archeal species and columns to (human) samples. Each entry corresponds to the number of non-human reads classified to the respective species. For convenience, in the following we refer to the set of microbial and viral species profiled using our approach as ‘metafeatures’.

By screening the study abstracts of the SRA for search terms prioritizing human clinical datasets derived from polyA-independent sequencing protocols (see Methods) we identified over 400 studies relevant to human disease comprising more than 17,000 cDNA libraries (close to 150 terabytes of raw sequencing data). Raw sequencing reads from these studies were downloaded and analyzed using the high performance computing system of the Leibniz Supercomputing Centre (LRZ) of the Bavarian Academy of Sciences and Humanities which facilitated ultra-fast processing with median speeds of 25 and 21 million reads per hour per core per run for the STAR and CLARK-S steps, respectively. Overall, of the total over 500 billion RNA-seq reads processed, around 91% could be mapped to the human genome. A fraction of 8.6% of all reads remained non-discriminative at the species level and defined as “unclassified”. 0.03%, 0.20% and 0.39% of all reads were assigned to archaeal, bacterial or viral metafeatures, respectively. Despite these relatively low percentages, the absolute numbers of reads classified were in the hundred millions to billions, enabling statistical analyses.

### Methods

#### High performance computing environment

Project computations including download, alignment of reads onto the human genome and metafeature quantification were made on the high performance Linux Cluster at the LRZ (www.lrz.de/services/compute/linux-cluster).

#### RNA-seq data retrieval

Raw next generation sequencing data were downloaded from the SRA. The R package *SRAdb* was downloaded on 23 May 2017 and used to query of the SRA database. To identify SRA projects that contain transcriptomic analyses of human RNA-seq data, the SRA attributes ‘taxon_id’, ‘library_source’, ‘library_strategy’, ‘platform’ were searched for the terms ‘9606’, ‘TRANSCRIPT’, ‘RNA-seq’, ‘ILLUMINA’, respectively. To remove potential bias derived from different sequencing technologies we also restricted the query to SRA runs annotated with ‘ILLUMINA’ in SRA attribute ‘platform’. To exclude studies with insufficient sample size for statistical analysis the query was restricted to SRA projects containing more than five runs. To avoid concentrating the analysis on a small number of large projects the query was restricted to SRA projects with less than 500 runs. To identify studies focusing on phenotypes relevant to human disease, we restricted the query to runs containing at least one or more of the terms ‘disease’, ‘patient’, ‘primary’ and ‘clinical’ in the SRA attribute ‘study_abstract’. To exclude *in vitro* (cell-culture) experiments, but focus on primary (clinical) samples, SRA runs containing the terms “mutant” or “cell-line” were removed from our selection. Furthermore, SRA runs containing the terms “single cell” and “GTEx” were removed. Finally, samples with less than 1 million total reads or read lengths <50 base pairs were excluded. The described query resulted in 484 Short Read Projects (SRPs) containing a total of 21,659 RNA-seq runs. Due to technical problems (i.e. missing URLs, restricted access) we were unable to download a fraction of 4,078 samples.

#### Human alignment

Alignment of reads against the human reference genome (hg38) and simultaneous human gene expression quantification was conducted with STAR (version 2.5.2). To increase mapping speed of a large number of samples, we used the --*genomeLoad LoadAndKeep* function to load the STAR index once and keep it in memory for subsequent alignments. The parameter --*quantmode GeneCounts* was used to generate the human gene expression count tables. Unmapped reads were saved with the -- *outReadsUnmapped Fastx* parameter. To further increase mapping speed, multiple threads were used as implemented with the parameter --*runThreadN 28*. Runs with less than 30 percent reads mapping to the human genome were excluded from downstream analysis. All human alignments were conducted on the LRZ “CoolMUC2” Linux-Cluster. This cluster contains 384 nodes with 64 GB RAM memory and 28 cores each.

#### Metafeature quantification

Metafeature quantification was conducted with CLARK-S (version 1.2.3).CLARK-S is a software method for fast and accurate sequence classification of metagenomic next-generation sequencing data, including RNA-seq data. One major issue during the classification of metagenomic data is the rising number of targets to align against. CLARK-S solves this issue by building a large index file consisting of discriminative *k*-mers. The metagenomic reference database was generated following the description of the CLARK website using the following two commands: 1) *set_targets.sh bacteria virus --species* and 2) *buildSpacedDB.sh*. This database contained a total of 16,551 genome sequences corresponding to 6,979 unique species (additional file 1). To allow uniform processing, paired-end sequencing experiments were analyzed independently. Each single unmapped reads file was used as input for CLARK-S with the following parameters: *classify_metagenome.sh --spaced-O* list of FASTQ files. To increase classification speed, the CLARK-S express mode was selected and multiple threads were used with parameters --*m 2* and *--n 32*, respectively. The output files of this step contain all input read identifiers with the corresponding metafeature classification. In the subsequent step, total counts are summarized for each feature with the *estimate_abundance.sh* command. To enable comparison across single-end and paired-end experiments, metafeature counts from paired-end experiments were averaged and subsequently rounded to conserve count distribution. To account for varying sequencing depths, metafeature abundance was estimated as the number of reads per million (RPM) total reads sequenced. Metafeature quantification was conducted on the LRZ “Teramem” Linux-Cluster. This cluster contains one node with 6,144 GB RAM memory and 96 cores.

#### BLAST based metafeature classification

To validate results generated by the MetaMap pipeline, the Basic Local Alignment Search Tool [24] was used as follows. A BLAST database was created from the same genome sequences used in the CLARK-S approach. Then, reads were aligned to this database using BLASTN with a threshold E-value of 1e-10. Produced counts from paired-end experiments were averaged. For each file, BLAST was done by running approximately 10 kilobase chunks (record separator “>") in parallel using GNU parallel (28 jobs), each with 8 threads using one node on the LRZ “CoolMUC3” Linux Cluster. This cluster contains 148 nodes with 96 GB RAM memory and 64 cores each. Output was parsed to exclusively keep reads that could be assigned at the species level.

#### Differential metafeature abundance

Differential metafeature abundance analysis was performed using the R package DESeq2 [25]. For each of the four published bona fide dual RNA-seq studies we classified samples into two groups based on the provided annotations: 1) Samples expected to contain the known pathogen, such as human papillomavirus positive head and neck tumors in the Zhang et al study, and 2) pathogen-free controls, such as mock-treated cells in the Westermann et al study. Using this binary outcome we performed differential expression analysis across all detected metafeatures. To account for sequencing depth, library size factors were estimated from the total number of sequenced reads. The dispersion for the negative binomial distribution was estimated using a local linear regression as implemented in the *DESeq()* function via the *fitType* parameter ‘local’.

### Data Validation and quality control

We validated our approach by recovering the ground truth in bona fide dual RNA-seq experiments performed with human cell lines and samples from patients with well-known infection status. Of the four selected studies, one analyzed an infection model based on a bacterial (*Salmonella enterica* serovar Typhimurium) and three based on distinct viral pathogens (Human papillomavirus, Herpes simplex virus, Rhinovirus). As expected, MetaMap detected the known pathogen at higher levels in the respective study compared to the other studies and pathogens (Table 1). Moreover, using the annotation provided in the respective study, we performed differential metafeature abundance analysis to identify those metafeatures that show the largest difference in abundance levels between the infected and control samples. The correct infection agent showed the most significant difference across all metafeatures between infected and control samples for each study (Fig. 2). For example, Westermann et al [26] generated dual RNA-seq data from HeLa cells infected with the enteric bacterial pathogen *Salmonella enterica* serovar Typhimurium and compared them to mock-treated control samples. Accordingly, we here observed *Salmonella enterica* as the most differentially abundant metafeature between the infected and the control samples (P<1e-75, Fig. 2A). Likewise we recovered *Alphapapillomavirus 9, Human alphaherpesvirus 1* (also known as herpes simplex virus 1) and *Rhinovirus A* as the most differentially abundant metafeatures in the data from Zhang et al [27], Rutkowski et al [28] and Bai et al [29], respectively. In the Westermann et al [26] and Rutkowski et al [28] studies, several additional metafeatures showed a strong differential abundance effect (Fig. 2A & C). These metafeatures were closely related to the true infection agent, i.e *Salmonella bongori* (P<1e-67) and *Panine alphaherpesvirus 3* (P<1e-9) for the Westermann et al [26] or Rutkowski et al [28] study, respectively. These findings confirm that our MetaMap pipeline recapitulates results from dedicated dual RNA-seq studies, i.e. studies based on known infectious agents. Therefore, MetaMap may be equally suited to detect previously unknown microbial and viral species in human primary samples.

**Table 1.**
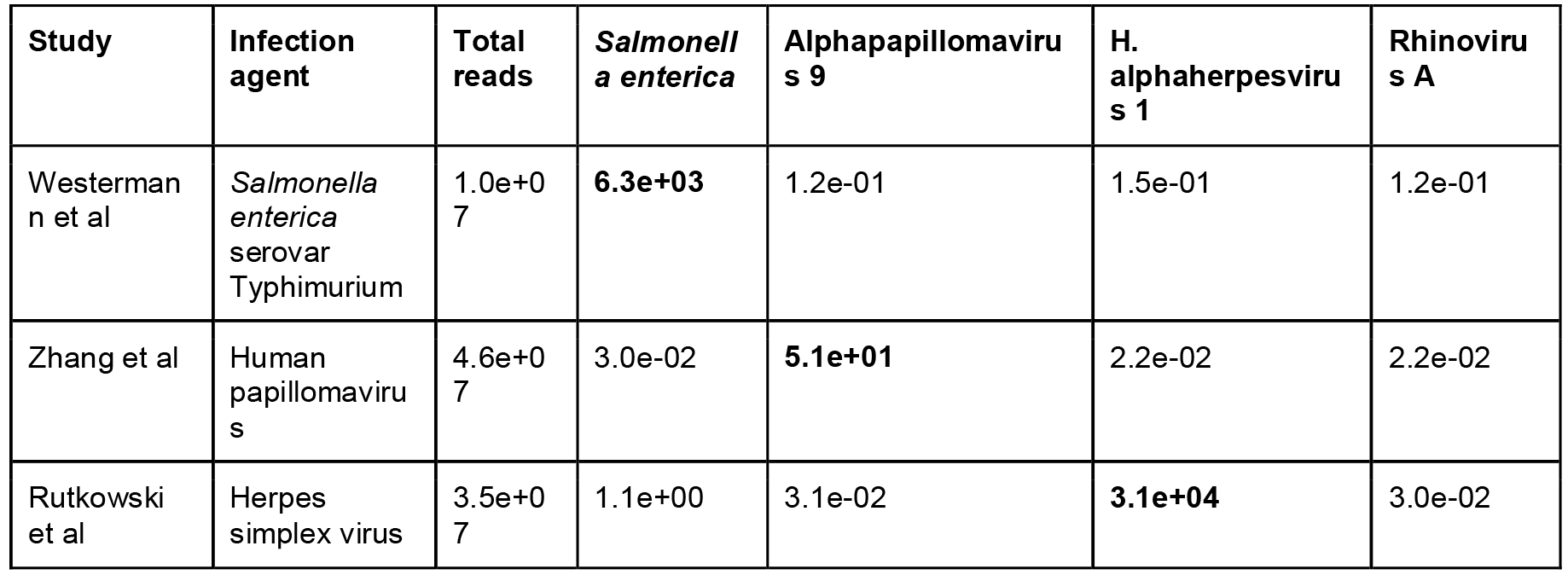
Overview of four dual RNA-seq studies used to validate the MetaMap pipeline. Total reads column depicts the average read depth per sample for each study. Average metafeature abundance for *Alphapapillomavirus 9*, *Salmonella enterica*, *Human alphaherpesvirus 1* and *Rhinovirus A* are shown in RPM. The correct infection agent for the respective study is highlighted in bold font.

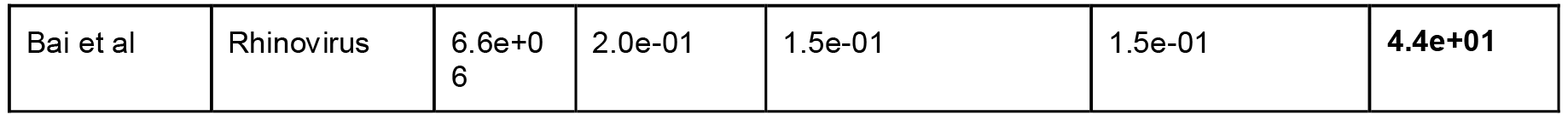

**Figure 2.**
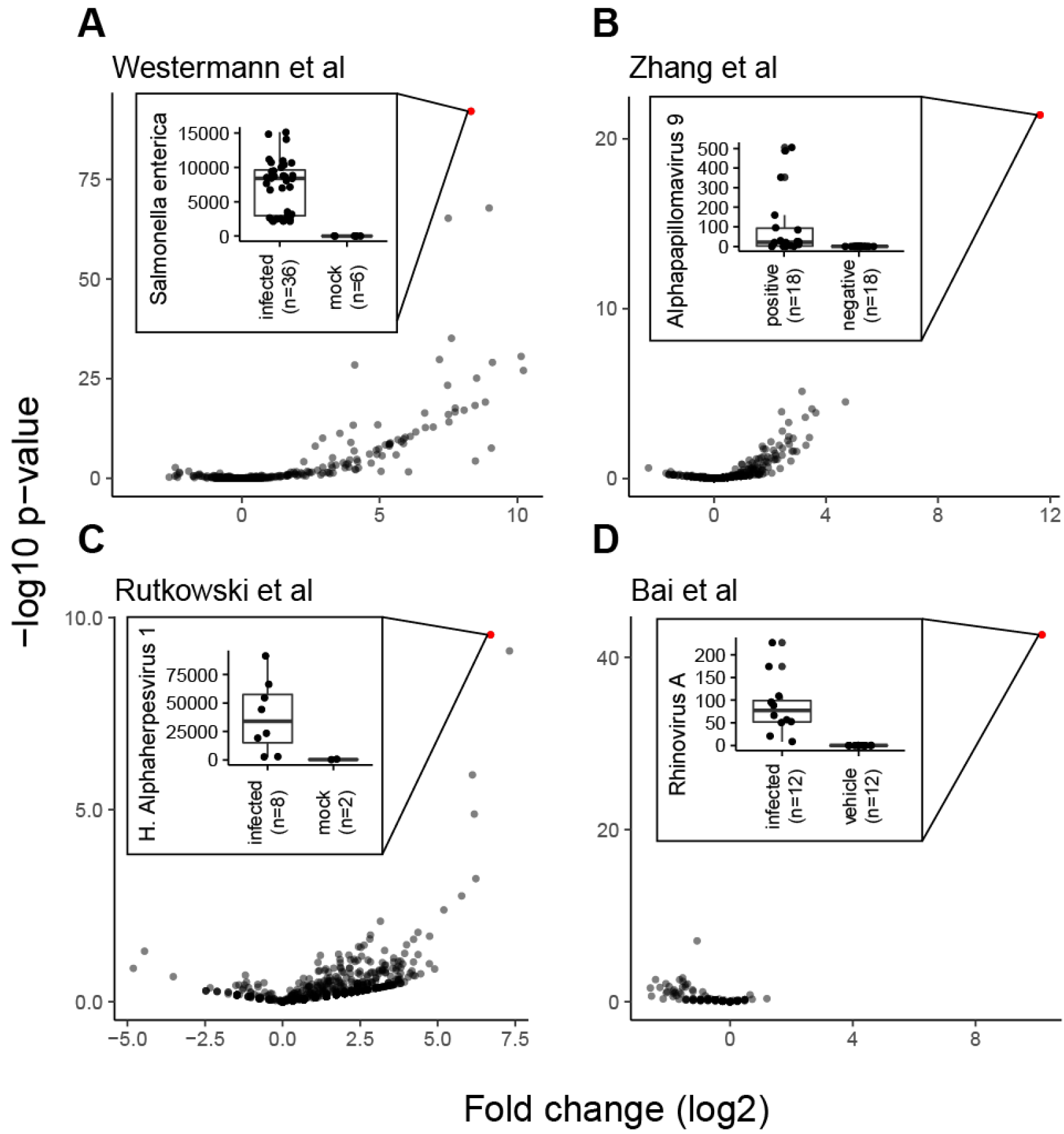
Differential metafeature abundance analysis of controlled infection experiments recovers ground truth. Panels A-D depict “volcano” plots showing fold change and inverted p-value on the X and Y axes, respectively. Each dot represents a metafeature. The most significant metafeature is colored in red. Insets display boxplots of the abundance levels in RPM of the top hit metafeature across conditions for each study. For all boxplots, the box represents the interquartile range, the horizontal line in the box is the median, and the whiskers represent 1.5 times the interquartile range.

As an additional control, we re-analysed two projects contained in our data collection that are derived from the B lymphoblast cell line, under non-infectious conditions. However, since Epstein-Barr virus is used for transfection and transformation of lymphocytes to lymphoblasts, we expected to detect reads from this virus in these projects [30], but no further viral or microbial reads [31]. Indeed the most abundant metafeatures in each project were dominated by reads classified to *Gammaherpesvirus 4* (also known as Epstein-Barr virus, EBV) and *Enterobacteria phage phiX174 sensu lato* (phiX), commonly used as spike-in in Illumina sequencing runs [32] (Fig. 3A-B). On average 95% and 97% of all metafeature reads were classified as phiX or EBV for projects SRP041338 and SRP091453, respectively (Fig. 3C). Conversely, the abundance of reads mapping to bacterial species for these two projects corresponds to the bottom percentile as compared to all other projects in the MetaMap database, supporting sterility of this cell line (Fig. 3D). This demonstrates that MetaMap not only is capable of re-discovering known pathogenic species (true positives) in controlled infection experiments (Fig. 2), but it also minimizes the detection of false positives or at least, provides measures such as abundance and significance allowing the user to identify and counterselect those species.

**Figure 3.**
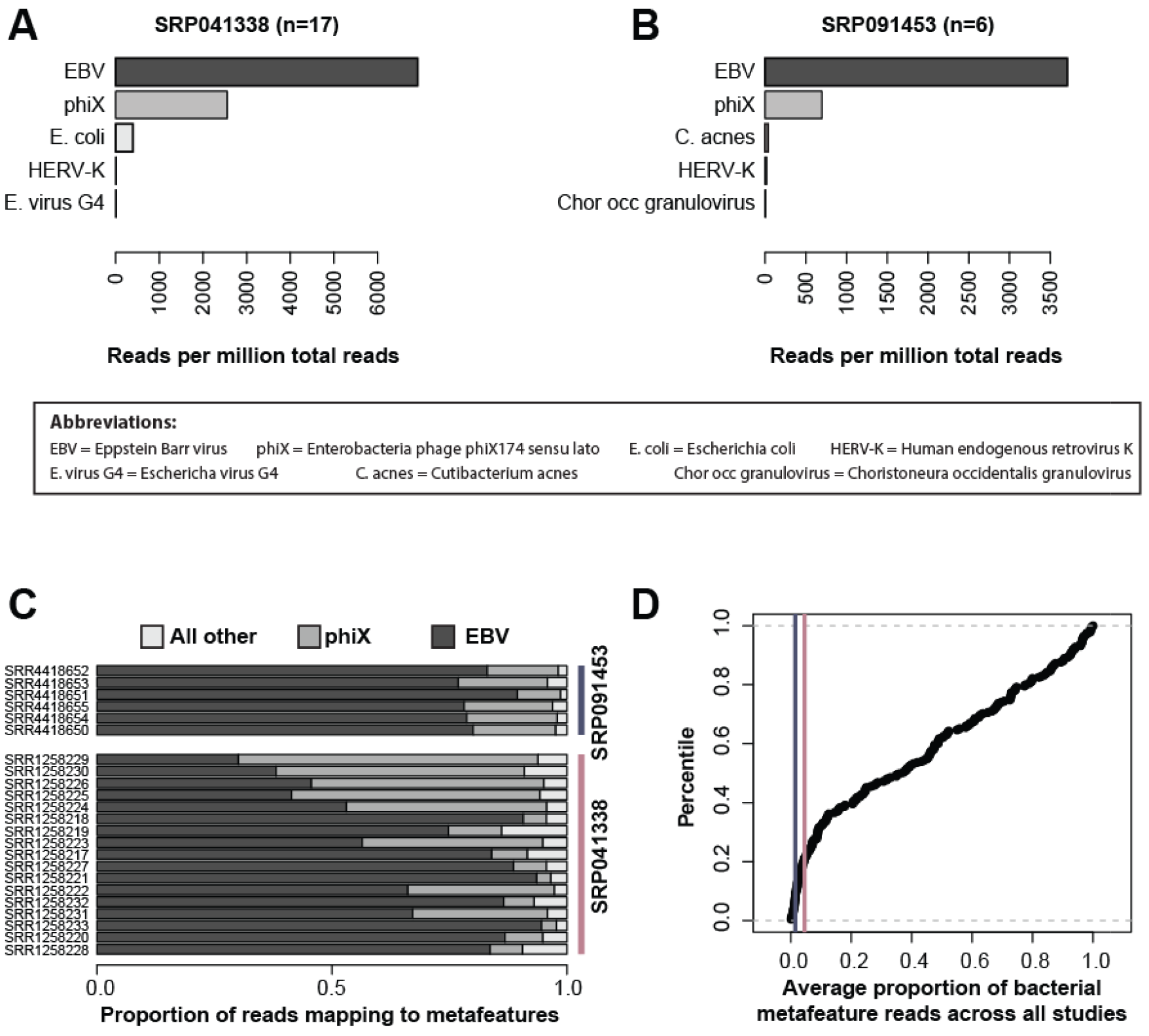
Analysis of lymphoblast cell line experiments further supports the MetaMap pipeline. Panels A and B depict mean abundance levels across all samples of the top five metafeatures for projects SRP041338 and SRP091453, respectively. Panel C shows relative proportion of reads mapping to EBV, phiX and all other metafeatures across RNA-seq samples. Panel D depicts the cumulative distribution plot of the average proportion of bacterial metafeature reads across all projects. Purple and pink vertical lines highlight projects SRP041338 and SRP091453, respectively.

As a technical validation, we compared our approach to an alternative metatranscriptomic classification strategy for the Westermann et al [33] study. All non-human reads were aligned using BLASTN to a BLAST database consisting of the same genomic sequences used by CLARK-S (see Methods for details). The average metafeature abundances across all 42 samples derived from the BLAST based approach and CLARK-S correlated significantly (Spearman correlation, Rho: 0.16, P: 3.1e-10) (Fig. 4A). BLAST showed higher sensitivity and detected more metafeatures compared to CLARK-S (indicated by the accumulation of dots at value 0 on the X-axis in Fig. 4A). This is mostly observed for low abundance metafeatures which could represent low counts derived from sequencing and/or mapping errors. However, most importantly the true pathogen metafeature ‘*Salmonella enterica*’ showed very high correlation across samples between the BLAST and CLARK-based abundance estimates (Fig. 4B). Noteworthy, the MetaMap pipeline processed reads more than three orders of magnitude faster than BLAST, demonstrating a significant speed advantage while generating comparable results (Fig. 4C).

**Figure 4.**
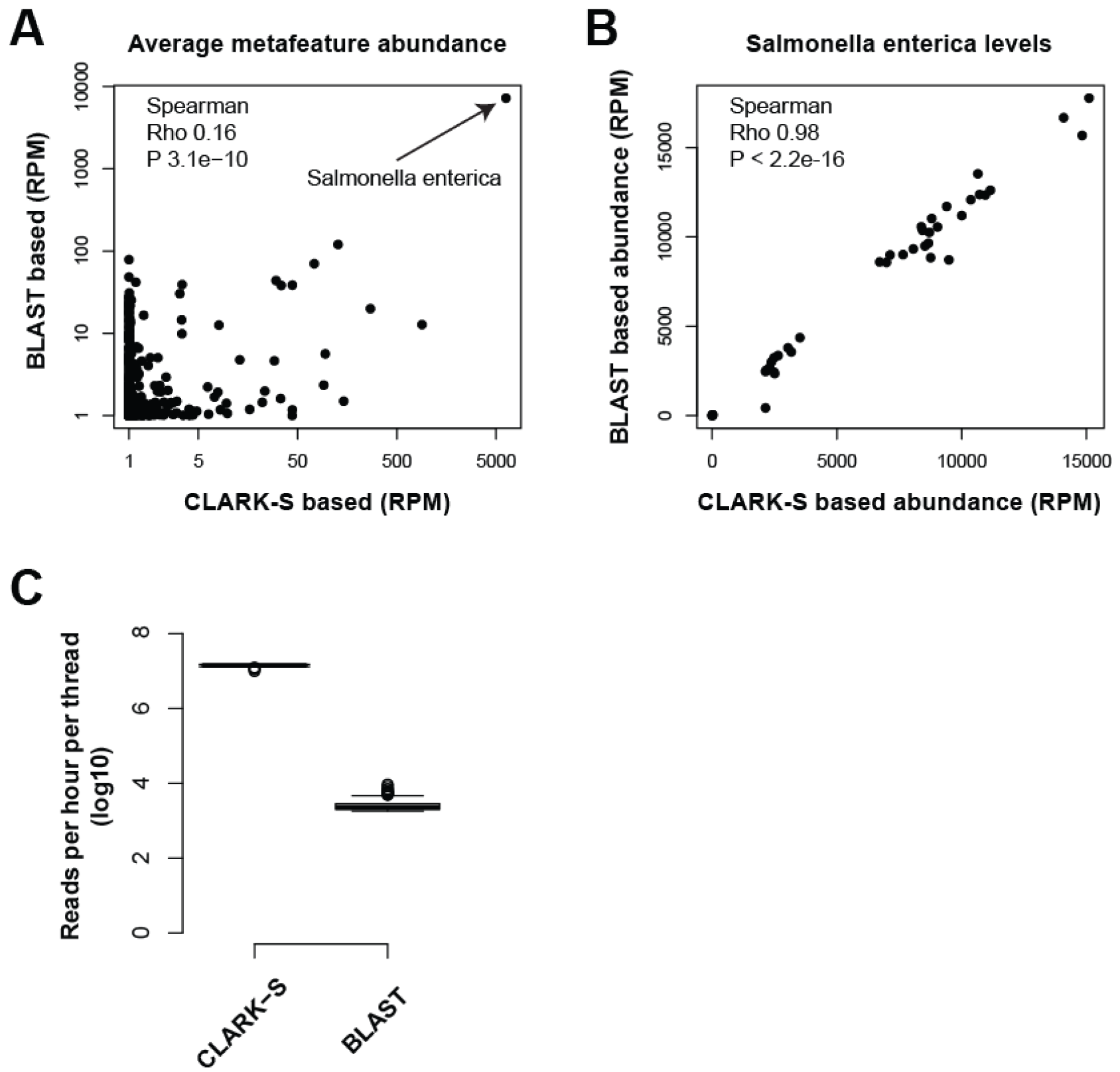
Alternative BLAST-based classification method validates metafeature abundance estimates by MetaMap. Panel A depicts average metafeature RPM levels derived using the CLARK-S software, as implemented in the MetaMap pipeline, and a BLAST-based alternative approach on the X- and Y-axes, respectively. Panel B shows the correlation in *Salmonella enterica* abundance levels between the two classification approaches. Panel C shows the difference in classification speed between the BLAST and CLARK-S metatranscriptomic classification. Y axis shows the number of reads processed per hour per thread in log10 space.

### Re-use potential

Microbial and viral contamination in next-generation sequencing data was observed before. It can be caused by incorrect mapping due to sequence similarity between different species [34,35]. To minimize such effects, we encourage focussing on studies including intra-project comparisons, such as exemplified in the differential metafeature abundance analysis. Contaminating agents should affect all runs within a project to the same extent and therefore not show a condition-specific effect. Alternatively, these “contaminations” might actually reflect true biological factors. For example, in the Westermann et al study [33] we detected substantial levels of phiX in both conditions (infected samples and mock-treated controls), but only the ‘*Salmonella*’ metafeature showed a condition-specific effect.

All the raw data described in the present study were publicly available before, yet have been very cumbersome to extract individually. The presented MetaMap database now makes these data easily accessible for a very broad community, thereby allowing for global comparisons over hundreds of individual studies and thousands of sampled conditions. While we attempted to minimize the risk of detecting false positives (Fig. 3), it should be noted that not all metafeatures classified by MetaMap will necessarily refer to true biological factors. Rather our pipeline provides the user with a scientific starting ground to validate the presence/absence of defined microbial and viral species under defined conditions and explore the underlying biology and significance in greater detail. As a potential use case of these data, users can test for associations of microbial or viral metafeatures with a plethora of human diseases, or between themselves. In addition, users with interest in a specific bacterial or viral species can easily identify studies, and consequently disease contexts, in which reads from this organism were detected. This could give an important first hint to assess whether the respective species might be implicated in a given human disease etiology. Furthermore, this resource provides the opportunity to validate findings derived from standard microbiome profiling technologies, such as 16S rRNA gene based or shotgun metagenomics [36]. Finally, metafeature detection in human clinical RNA-seq samples may provide a critical advantage when studying microbes or viruses which are challenging to isolate.

All generated metafeature OTU count tables from 17,278 cDNA libraries from 436 SRA projects including annotation are provided for download. The MetaMap pipeline can be accessed via the protocols.io website with digital object identifier dx.doi.org/10.17504/protocols.io.msec6be.

## Acknowledgments

The authors would like to thank Yu Wang and Ferdinand Jamitzky from the LRZ for their support.

